# Axonal injury signaling is restrained by a spared synaptic branch

**DOI:** 10.1101/2024.11.07.622420

**Authors:** Laura J. Smithson, Juliana Zang, Lucas Junginger, Thomas J. Waller, Reilly Jankowiak, Sophia Khan, Ye Li, Dawen Cai, Catherine A. Collins

## Abstract

The intrinsic ability of injured neurons to degenerate and regenerate their axons facilitates nervous system repair, however this ability is not engaged in all neurons and injury locations. Here we investigate the regulation of a conserved axonal injury response pathway with respect to the location of damage in branched motoneuron axons in *Drosophila* larvae. The dileucine zipper kinase DLK, (also known as MAP3K12 in mammals and Wallenda (Wnd) in *Drosophila*), is a key regulator of diverse responses to axonal injury. In three different populations of motoneurons, we observed the same striking result that Wnd/DLK signaling becomes activated only in response to injuries that remove all synaptic terminals. Injuries that spare even a small part of a synaptic terminal fail to activate Wnd/DLK signaling, despite the presence of extensive axonal degeneration. The regulation of injury-induced Wnd/DLK signaling occurs independently of its previously known regulator, the Hiw/PHR ubiquitin ligase. We propose that Wnd/DLK signaling regulation is linked to the trafficking of a synapse-to-nucleus axonal cargo and that this mechanism enables neurons to respond to impairments in synaptic connectivity.

## Introduction

Repair of nervous system damage requires an ability of neurons to regenerate axons and synaptic connections. However this ability is not universally induced following injury, most notably following spinal cord injury (SCI). While many studies have identified extrinsic factors that influence or inhibit axonal regeneration, several studies have noted that the location of damage can influence the intrinsic ability of the neuron to mount a transcriptional response to the damage (Fernandes et al., 1999; Lorenzana et al., 2015; Mason et al., 2003; Wang et al., 2024). Some studies have noted an effect of distance from the cell body for long axons in the spinal cord (Fernandes et al., 1999; Mason et al., 2003; Wang et al., 2024). Other studies have noted that the location of injury with respect to an axonal branching point also strongly influences the response (Lorenzana et al., 2015; Wu et al., 2007). Even in a strongly inhibitory environment to regeneration, dorsal column sensory axons show robust axonal growth when injured proximal to their bifurcation in the spinal cord (Lorenzana et al., 2015).

Here we investigate the regulation of a conserved axonal injury response pathway with respect to the location of axonal injury. The dileucine zipper kinase DLK (also known as MAP3K12 in mammals and Wallenda (Wnd) in *Drosophila*), is a key regulator of diverse responses to axonal injury. These include an essential role in the ability of damaged neurons to initiate axonal regeneration in worm and fly models (Chen et al., 2011; Hammarlund et al., 2009; Stone et al., 2014; X. Xiong et al., 2010; Yan et al., 2009), synaptic repair and recovery following CCR5 inhibition in a stroke model (Joy et al., 2019), and enhanced regeneration and mechanical allodynia following PNS nerve damage in mice(Hu et al., 2019; Wlaschin et al., 2018). Dichotomously, DLK is also required for the death of retinal ganglion cells following optic nerve damage (Watkins et al., 2013; Welsbie et al., 2017, 2013). In mammalian as well as in fly neurons, this kinase associates with vesicles that are physically transported in axons (Holland et al., 2016; Xiong et al., 2010), while downstream nuclear signaling requires functional axonal transport machinery (Xiong et al., 2010). DLK is therefore considered to function as a ‘sensor’ of axonal damage, whose activation can confer responses of repair or death, depending upon the cellular context(Asghari Adib et al., 2018).

While the responses gated by DLK are impactful for neurons and their circuits, the mechanism(s) that lead to DLK signaling activation are still poorly understood. A number of observations have documented DLK signaling activation in neurons that are not mechanically damaged but have experienced some form of cellular stress. These include the presence of cytoskeletal mutations (Bounoutas et al., 2011; Chen et al., 2014; Kurup et al., 2015; Valakh et al., 2013) and the presence of chemotherapy agents (Bhattacharya et al., 2012; DeVault et al., 2024; Valakh et al., 2015) known to impair axonal cytoskeleton integrity and transport. DLK activation is also responsible for phenotypes associated with mutations in the unc-104/KIF1A kinesin (Li et al., 2017), a major carrier of synaptic vesicle precursors in axons (Guedes-Dias and Holzbaur, 2019; Petzoldt, 2023). Inhibition of DLK is protective in mouse models of ALS and Alzheimer’s Disease (Le Pichon et al., 2017; Patel et al., 2017). These observations have fostered growing interest in DLK as a potential therapeutic target, and in understanding the mechanisms that control DLK signaling activation in neurons.

Here we test the hypothesis that DLK/Wnd signaling is tuned to the synaptic connectivity of a neuron. A shared feature of nerve injuries and stressors that disrupt axonal cytoskeleton and transport is a loss in downstream connections. A conserved regulator of DLK/Wnd, the E3-ubiquitin ligase PAM/Highwire/Rpm-1/Phr1 (Collins et al., 2006; Huntwork-Rodriguez et al., 2013; Nakata et al., 2005), is hypothesized to function at synaptic terminals (Abrams et al., 2008; Opperman and Grill, 2014; Xiong et al., 2012; Zhen et al., 2000). This led us to ask whether interactions at an intact synaptic terminal are responsible for restraining Wnd signaling in uninjured neurons. We probed this hypothesis through injuries to branched motoneuron axons in *Drosophila* larvae, which allowed us to compare injuries that leave spared synaptic terminals to injuries that lead to complete denervation. In three different populations of motoneurons, we observed the same striking result that Wnd signaling becomes activated only in response to injuries that remove all synaptic terminals. Injuries that spared even a small part of a synaptic terminal did not activate Wnd signaling, despite the presence of extensive axonal degeneration. Surprisingly, removal of all synapses led to additive induction of Wnd signaling in *hiw* mutants. These observations suggest the existence of a mechanism that restrains Wnd signaling at synaptic terminals independently of the Hiw ubiquitin ligase.

## Results

### The presence of a spared synaptic branch restrains Wnd-mediated injury signaling in SNc motoneurons

To determine whether synaptic connections influence injury signaling by Wnd/Dlk, we established methods to injure single synaptic branches of defined larval motoneurons. The m12 (5053A)-Gal4 driver line (Ritzenthaler et al., 2000) that we have used in previous nerve injury studies (Xiong et al., 2010; Xiong and Collins, 2012), drives expression in two single motoneurons that project closely fasciculated axons to innervate body wall muscles 26, 27 and 29 (**Figure 1A**). This pattern was previously attributed to a motoneuron (MN) named MNSNc, which was noted to have ‘two cell bodies’ (Kim et al., 2009). We used the Bitbow2 (Li et al., 2021) multi-colored cell labeling approach to resolve the two neurons, and observed an invariable pattern that one MNSNc neuron innervates muscles 26 and 29, while the other innervates muscle 27. (**Figure 1B** shows two examples: the neuron that innervates muscle 27 (middle images) expresses a set of colors that distinguish it from the terminals on muscles 26 and 29). We therefore refer to these two neurons labeled by the m12Gal4 driver as MNSNc-26/29 and MNSNc-27 (**Figures 1A, B, Supplemental Figures 1A**). MNSNc-26/29 (cartooned in blue in **Figure 1A**) has three collateral branches: two branches innervate muscle 26, and a single branch innervates muscle 29. MNSNc-27 (cartooned in red in **Figure 1A**) has two collateral branches that innervate muscle 27: these two branches most often remain together and occasionally bifurcate separately onto muscle 27 (**Supplemental Figure 1A**). This innervation is stereotyped across segments and animals. As previously noted (Kim et al., 2009), the paired cell bodies of the MNSNc neurons are found on the lateral sides of the abdominal region of the ventral nerve cord (VNC). Bitbow2 expression revealed that the MNSNc-27 cell somas are positioned more ventrally compared to the cell somas of MNSNc-26/29 in the VNC (**Supplemental Figure 1B**).

**Figure 1:**
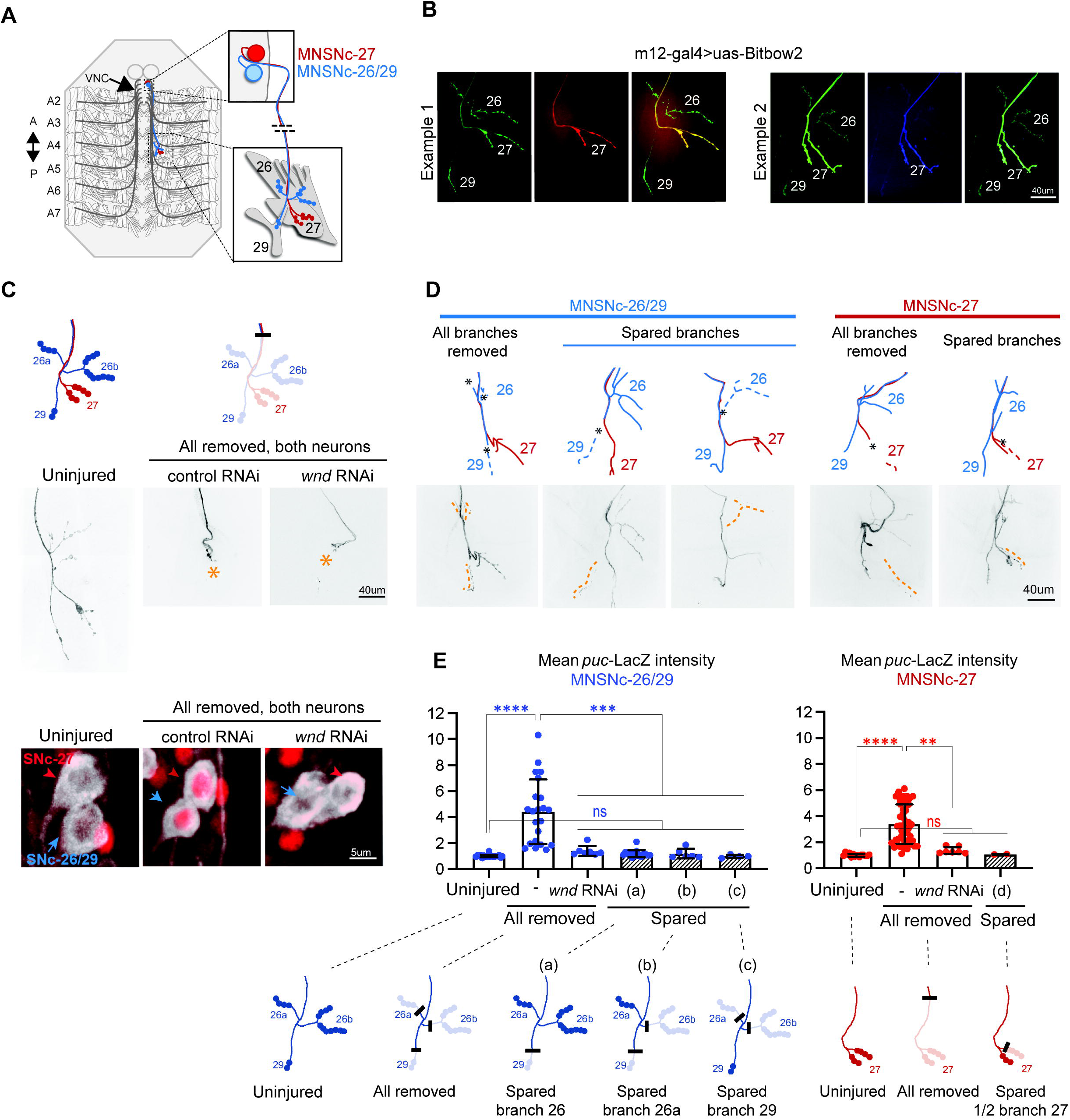
A spared synaptic branch restrains Wnd-dependent injury signaling in SNc motoneurons. **A)** Schematic representation of the two SNc motoneurons innervating muscles 26 and 29 (MNSNc-26/29, red) and muscle 27 (MNSNc-27, blue), which are labeled by expression of the m12-Gal4 driver. **B)** Example images of NMJ terminals from m12-Gal4/+; UAS-BitBow2 (Li et al., 2021) /+ third instar larvae, used to define the connectivity shown in A. The neuron that innervates muscle 27 (MNSNc-27) expresses a distinct set of colors from the Bitbow2 (Li et al., 2021) reporter than the neuron that innervates muscles 26 and 29 (MNSNc-26/29). **C)** Confirmation of *puc*-lacZ induction following laser axotomy. The cartoons on the top row show the location used to injure both axons; this location removes all of the synaptic branches from both MNSNc-26/29 and MNSNc-27. The middle row shows example injuries (versus uninjured, right) at the indicated location in m12-Gal4, UAS-mCD8GFP/*puc*-lacZ larvae. The bottom row shows examples of *puc*-lacZ expression (red channel) in the MNSNc cell bodies 24h following injury. **D)** Example MNSNc-26/29 (blue) and MNSNc-27 (red) neurons injured at different locations. **E)** Quantification of *puc*-lacZ intensity measurements in MNSNc-26/29 (blue) and MNSNc-27(red) following injuries that remove all synaptic branches versus injuries that leave a spared synaptic branch. Injury location (a) removes the small number of boutons on muscle 29 while sparing the boutons on muscle 26. Injury location (b) removes boutons from muscle 29 and the posterior sub-branch on muscle 26. Injury location (c) removes all branches except for the small number of boutons on muscle 29. Note that all injuries that leave spared boutons (hatched shading) show no *puc*-lacZ induction, regardless of the number of boutons lost or spared. A one-way ANOVA with Tukey test for multiple comparisons was performed for each neuron. **** p < 0.0001; ** p = 0.0011; ns = not significant.t.

We used a pulsed dye laser to carry out axotomies of MNSNc motoneuron axons at different locations in intact immobilized larvae (described in methods and (Smithson et al., 2024a)). Successful injuries were confirmed by the degeneration of distal stumps within 24h post-injury. To assess the activation of Wnd signaling, we probed for the induction of *puckered* expression from the *puc*-lacZ enhancer trap (Martín-Blanco et al., 1998). Previous studies from this reporter line have shown that expression of nuclear localized lacZ is strongly induced following nerve injury, requiring Wnd, JNK and the Fos transcription factor (Xiong et al., 2010). We confirmed this is the case for MNSNc neurons; axotomy of the m12-Gal4 labeled neurons upstream of the synaptic branches led to a four fold induction of *puc*-lacZ expression in either neuron 24h after the injury; this was abolished in neurons that co-express double stranded RNAi targeting *wnd* (**Figure 1C**). Similar results were observed for nerve crush injuries (**Supplemental Figure 1C**).

In contrast to axotomies that removed all synaptic branches, laser injury to single collateral branches of MNSNc-26/29, including branches innervating either muscle 26 or 29 and resulting in spared synaptic branches, fail to induce *puc*-lacZ expression (**Figures 1D, E)**. In addition, laser ablation of the anterior branch of MNSNc-27 also failed to activate Wnd signaling (**Figures 1D, E**). We note that the induction of puc-lacZ did not correlate with the number of boutons that were lost or spared. MNSNc-26/29 forms four-fold more boutons on muscle 26 (17 +/- 3.8) than 29 (4 +/- 1.3). However injuries that spared any of the branches, even the small number on muscle 29, showed equivalent puc-lacZ levels to uninjured neurons (**Figure 1E**; compare injury locations a, b and c). We carried out similar experiments in aCC motoneurons, which can be labeled with the Dpr4-Gal4 driver) (Pérez-Moreno and O’Kane, 2019)). Laser axotomies that removed all of the synaptic branches resulted in four-fold increase in *puc*-LacZ levels, while injuries that left spared synaptic boutons failed to induce *puc*-lacZ expression. (**Supplemental Figure 1D,E**). These observations suggested that even a small number of remaining boutons was sufficient to restrain the activation of *puc*-lacZ expression.

Despite the small distance from the disconnected muscle, none of the injured MNSNc synaptic branches were able to re-innervate the muscle. We think this is due to an absence of axon growth-promoting cues, since MNSNc axons did show robust but misdirected axonal growth into the segmental nerve SNa following injuries in locations upstream of the synaptic branches (data not shown). However we did notice differences in the trafficking of proteins to injured proximal stumps. An example of this is shown for ectopically expressed kinase-dead Wnd transgenic protein, GFP-Wnd-KD in **Supplemental Figure 1F, G.** (We were only able to track kinase-inactive Wnd since overexpression of Wnd causes dramatic morphological defects to neurons (Collins et al., 2006; Feoktistov and Herman, 2016; Xiong et al., 2010). GFP-Wnd-KD was strongly induced and accumulated at the proximal stump following injuries that removed all synaptic branches, but was barely detectable following injuries that left spared synaptic branches (**Supplemental Figure 1F, G**). Collectively, these observations suggest that the presence of spared synaptic branches affect the subsequent events that occur in the injured axon. These include the stability and/or trafficking of Wnd protein in injured axons and the activation of Wnd-regulated signaling in the neuron soma.

### Restraint of Wnd-dependent injury signaling in bifurcated axons of type II VUM motoneurons

A more extreme example of axonal branching is illustrated by the ventral unpaired median (VUM) neurons, which project symmetric bifurcated axons through separate nerves to innervate multiple body wall muscles on both left and right sides of the larva (**Figure 2A**, **Supplemental Figure 2** and (Koon et al., 2011; Monastirioti et al., 1995; Vömel and Wegener, 2008). VUM neurons can be specifically labeled based on their expression of tyrosine decarboxylase 2 using the Tdc2-Gal4 driver. Each abdominal segment has 3 VUM neurons, each of which sends a single axon dorsally which then bifurcates in the midline VNC (**Supplemental Figure 2,** and (Vömel and Wegener, 2008). The two bifurcations then project through separate nerves to symmetrically innervate both left and right halves of the larval body. Through nerve crush injuries to only one ventral side of the animal, we were able to injure one bifurcation while leaving the other bifurcation intact. Successful injuries were determined based on the degeneration of the distal axon and synaptic terminals at 24 hours post-crush (**Figure 2B**). Whether both bifurcations, a single bifurcation or neither bifurcation was injured was scored for each VUM neuron by tracing the Tdc2-Gal4, UAS-mCD8GFP labeled axons from the nerve to the cell body.

**Figure 2:**
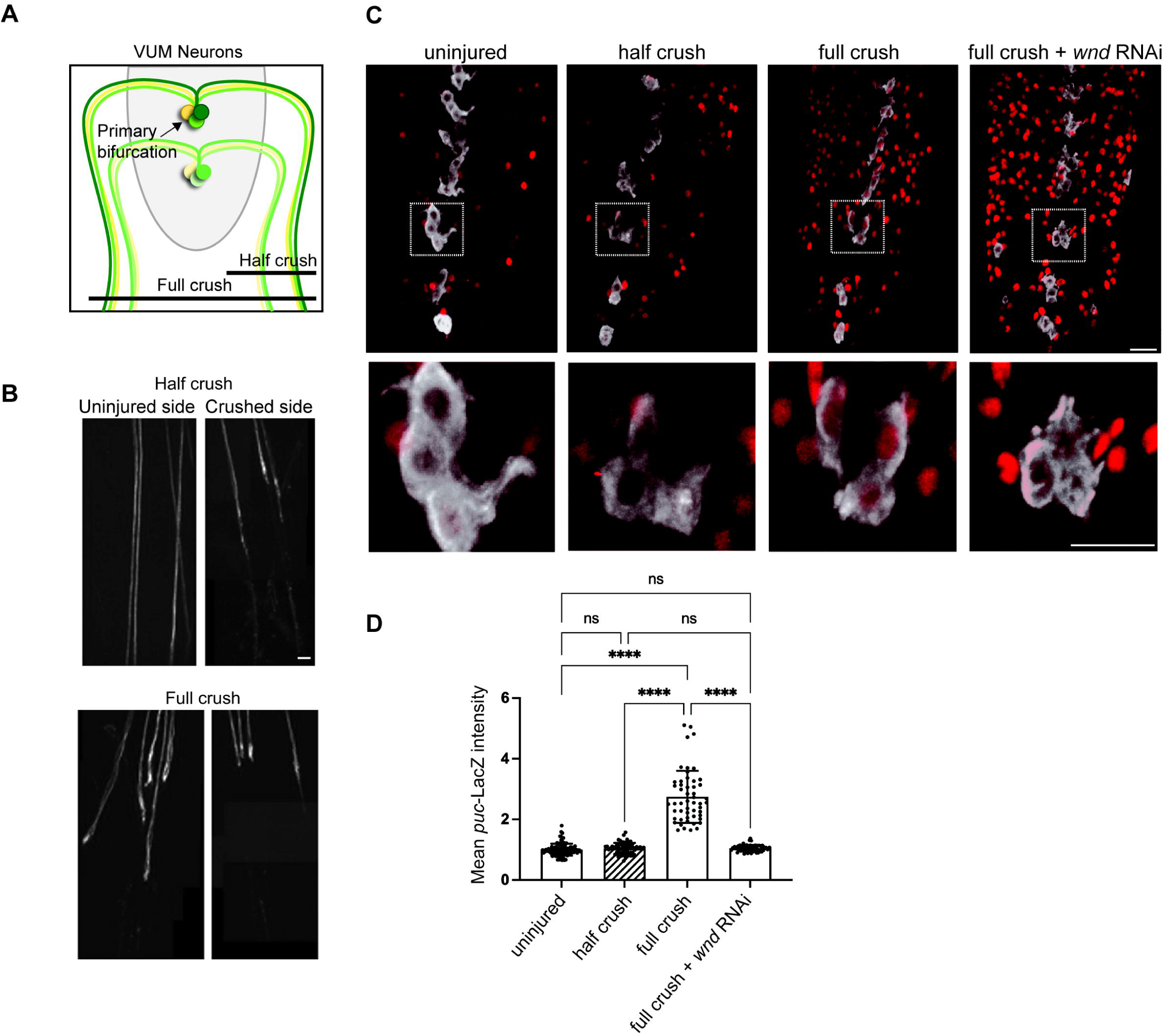
Restraint of Wnd-mediated injury signaling by spared branch in bifurcated neurons. **A)** Cartoon of Ventral Unpaired Motoneurons (VUM), which have bifurcated axons that symmetrically innervate body wall muscles on both the left and right sides of the animal. Nerve crush to either left or right side of the animal can axotomize a single bifurcation while leaving the other bifurcated axon intact. **B)** Example images of VUM axons (visualized in Tdc2-Gal4, UAS-mCD8-GFP larvae) in segmental nerves on the uninjured and injured sides following nerve crush to a single side. **C)** Example images of *puc*-lacZ expression in the VNC (ventral nerve cord) of larvae following nerve crush to a single side (half crush) versus crush to all the segmental nerves (full crush). *puc*-lacZ expression (red) is induced in VUM neurons (white) only after full crush. In contrast, other motoneurons, which innervate a single side, are induced by both half and full crush injuries. Co-expression of UAS-*wnd*-RNAi in VUM neurons cell autonomously inhibits *puc*-lacZ induction. **D)** Quantification of *puc*-lacZ intensity measurements in VUM neurons. A one-way ANOVA with Tukey test for multiple comparisons was performed. **** p < 0.0001; ns = not signifcant. Scale bars = 20 um.

Similarly to other motoneurons (Xiong et al., 2010), *puc*-lacZ expression is barely detectable in uninjured VUM motoneurons and, compared to uninjured VUM neurons, is induced almost 3-fold at 24h following complete/full nerve crush injuries that damage both bifurcations and remove all synaptic connections (**Figure 2C,D**). This induction is abolished in VUM neurons that co-express *wnd*-RNAi (**Figure 2C,D**). Note that *wnd*-RNAi is only expressed in the VUM neurons and does not affect the other MNs which do not express Tdc2-Gal4. In contrast to full nerve crushes, injuries to nerves on a single side of the animal that the other bifurcation intact (‘half crush’ injuries) failed to induce *puc*-lacZ expression in VUM neurons (**Figure 2C,D**). Most MNs make ipsilateral and not bilateral projections; in ‘half-crush’ injuries, *puc*-lacZ is induced in most non-VUM motoneurons on the side of the crush, but is not induced in VUMs. These combined observations suggest that in *Drosophila* motoneurons, Wnd signaling is not tuned to detect axonal damage *per se*, but is instead uniquely tuned to detect a complete loss of innervation, which occurs following some injuries but not others.

### Restraint of Wnd signaling at spared branches does not require synaptic transmission

Since the presence of intact synaptic boutons restrains Wnd signaling activation, we considered whether cellular events associated with evoked or spontaneous synaptic transmission are associated with this mechanism. Summarized in **Supplementary Table 1**, we tested a total of 22 genetic manipulations expected to inhibit synaptic transmission, but none led to a change in *puc*-lacZ expression. These include electrical silencing of SNc motoneurons by Gal4/UAS mediated expression of the *Drosophila* open rectifier K+ channel (dORK) (Nitabach et al., 2002), silencing of transmission using temperature-sensitive mutations in dynamin (Kitamoto, 2001), and light-induced silencing of neurons expressing *Guillardia theta* anion channelrhodopsin 1 (gtACR1)(Mohammad et al., 2017) (**Supplementary Table 1**). Consistent with these negative results, we noted that previous studies have described many genetic manipulations that perturb evoked and/or spontaneous synaptic transmission (Choi et al., 2014; Daniels et al., 2006; DiAntonio and Schwarz, 1994; Han et al., 2022) do not yield synaptic phenotypes (of synaptic overgrowth or decreased VGlut expression levels) associated with Wnd activation (Collins et al., 2006; Li et al., 2017).

### Restraint of Wnd-mediated axon injury signaling is independent of the Highwire ubiquitin ligase

We then asked whether a known upstream regulator of Wnd signaling, the Pam/Hiw/Rpm-1 (PHR) ubiquitin ligase, functions to restrain Wnd at synaptic branches. PHR is a large protein with multiple evolutionarily conserved domains, inducing a RING-finger domain, which regulates DLK/Wnd in invertebrate (*C. elegans* and *Drosophila*)(Collins et al., 2006; Nakata et al., 2005) and vertebrate (Huntwork-Rodriguez et al., 2013) model organisms. PHR is an attractive candidate since it localizes to synaptic terminals (Abrams et al., 2008; Opperman and Grill, 2014; Zhen et al., 2000) and loss of PHR function leads to increased levels of Wnd/DLK at synapses (Collins et al., 2006; Nakata et al., 2005). This hypothesis predicts that removal of all synaptic branches would be equivalent to a genetic loss in PHR function. We tested this in null mutants for the *Drosophila* ortholog of PHR, Highwire (Hiw). The *hiw*^Δ*N*^ mutation deletes the entire N-terminal half of the *hiw* gene and abolishes expression of the Hiw protein (Wu et al., 2005). Since *hiw*^Δ*N*^ animals are viable, we were able to carry out injury assays in neurons that completely lack Hiw function.

Consistent with previously reported phenotypes for *hiw* in other motoneuron types (Collins et al., 2006; Wan et al., 2000; Wu et al., 2005), uninjured MNSNc neurons in male *hiw*^Δ*N*^ mutants have an increased number of axon collateral and terminal branches at muscles 26, 29 and 27 NMJs (**Figure 3A** and **Supplemental Figure 3A**). Also consistent with previous observations, uninjured neurons show elevated expression of *puc*-lacZ in *hiw*^Δ*N*^ mutants compared to control animals (**Figure 3** and **Supplemental Figure 3A**). Strikingly, injuries that removed all synaptic terminals led to an even further elevation of *puc*-lacZ expression in *hiw*^Δ*N*^ mutant neurons. This was the case for laser axotomies that removed all synaptic branches from either MNSNc-26/29 and/or MNSNc-27 neurons (**Figure 3A**, right column, and **Figure 3B**), and also for VUM neurons following nerve crush injuries (Figure 3C). Injuries to a single synaptic branch (on muscle 29) of MNSNc-26/29 in *hiw*^Δ*N*^ mutants had a similar level of *puc*-lacZ expression as uninjured *hiw*^Δ*N*^ neurons (**Figures 3A, B**). Collectively, these observations suggest that the presence of a spared synapse is capable of restraining Wnd signaling independently of Hiw’s function.

**Figure 3:**
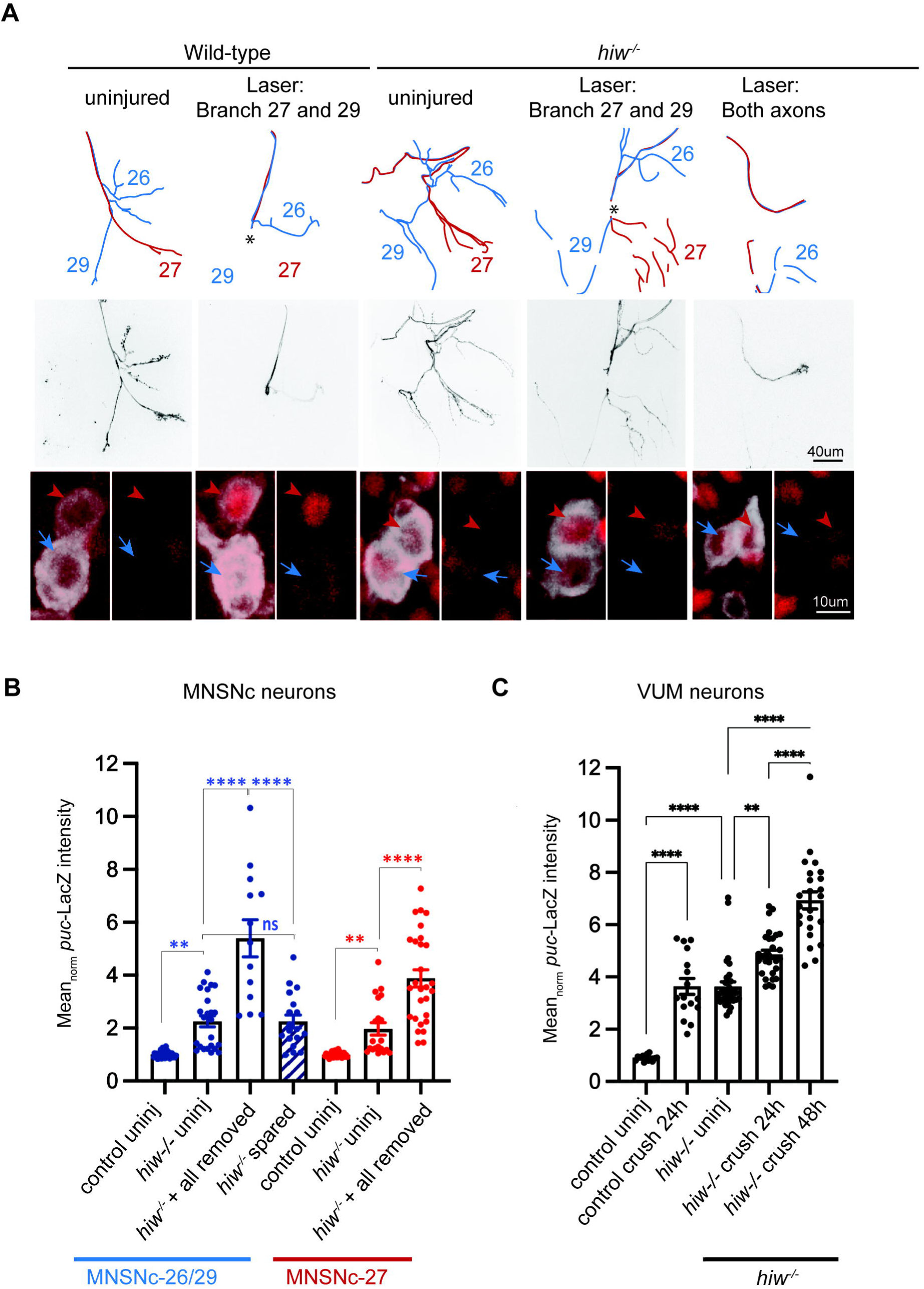
Presence of spared synaptic branch restrains Wnd signaling independently of Hiw. **A)** Laser axotomy is carried out to MNSNc neurons at a location that completely removes the synaptic terminal of MNSNc-27 (red neuron). The injury also leads to loss of the MNSNc-26/29 (blue) terminal on muscle 29 but not 26 (hence leaves a spared synaptic branch. The final column shows an axotomy that fully removes the terminals for both MNSNc neurons. These injuries were repeated in control animals versus the background of a *hiw* null mutant, *hiw*^Δ*N*^. **B)** Quantification of *puc*-lacZ expression for individual MNSNc neurons after full versus spared axotomies, compared to uninjured neurons. Basal *puc*-lacZ expression is already elevated in uninjured *hiw*^Δ*N*^ neurons compared to control. This can be further elevated in axotomies that remove all synapses, but not in axotomies that leave spared branches. **C)** Quantification of *puc*-lacZ in VUM neurons (labeled by Tdc-2-Gal4; UAS-mCD8-GFP) 24h and 48h following full nerve crush in control versus *hiw*^Δ*N*^ mutants. A two-way ANOVA with Tukey test for multiple comparisons was performed. **** p < 0.0001; *** p < 0.001; **p<0.01; ns = not significant.

## Discussion

### Axonal branches and spared synaptic connections influence the ability of injured axons to regenerate

Since the foundational observations of Ramon y Cajal, we have known that the ability of axons to regrow following damage varies strongly according to the location of damage (Ramón y Cajal, 1928). Most widely considered are differences in extrinsic factors that influence the ability of axons to grow following PNS injuries, and the impediments to axonal regeneration in the CNS that inhibit repair following spinal cord injuries (Huebner and Strittmatter, 2009). A less well studied but important intrinsic determinant of regeneration ability is the location of the injury with respect to axonal branches. For *C. elegans* PLM mechanosensory neurons, robust regeneration occurs following injuries that remove both the synaptic and sensory branches, but not following injuries that leave a synaptic branch intact (Wu et al., 2007). In the mammalian spinal cord, an elegant study following responses to laser-induced microsurgery to ascending and descending central projections of sensory neurons observed that remarkable regeneration occurred following injuries proximal to the branching point (which led to removal of all branches) but not distal (which left one branch intact) (Lorenzana et al., 2015). Studies in mice have also implicated synaptic proteins alpha2-delta, Munc13 and RIM in restraining regenerative ability of axons (Hilton et al., 2022; Tedeschi et al., 2016). These observations are consistent with the possibility that spared afferent synaptic connections that remain following injuries distal to the branch point inhibit the regeneration ability of centrally projecting axons in the spinal cord.

It is noteworthy that many of the axons that project over great distances in the spinal cord have at least one synaptic branch. Neurons that project through the corticospinal tract (CST), whose poor regeneration ability following spinal cord injury is most widely studied, form synaptic branches in the red nucleus, brainstem, and throughout the spinal cord (Raineteau and Schwab, 2001). A recent study has profiled the responses of CST neurons at different injury locations (Wang et al., 2024). Strikingly, regeneration associated genes (RAGs) and phosphorylated cJun, a marker of DLK signaling activation, are not induced in CST neurons following spinal cord injury, but are robustly induced following intracortical injuries close to CST cell bodies (Mason et al., 2003; Wang et al., 2024). Since the latter may be the only form of injury that removes all synaptic branches from the CST neurons, we propose that restraint of DLK signaling activation by spared synaptic branches could be a prominent feature of the poor intrinsic regeneration capacity of neurons following spinal cord injury.

### Restraint of Wnd/DLK signaling at synaptic terminals

Using genetic manipulations that inhibit or perturb synaptic transmission and/or neuronal excitability, we did not detect a requirement for synaptic transmission in the restraint of Wnd by spared synaptic branches. The PHR ubiquitin ligase, known as Hiw in *Drosophila*, was a logical candidate to regulate Wnd at synapses, since studies in multiple model organisms have shown that loss of this enzyme leads to increased levels of Wnd/DLK in axons (Collins et al., 2006; Huntwork-Rodriguez et al., 2013; Nakata et al., 2005). The *puc*-lacZ reporter implies that Wnd signaling is elevated in *hiw* mutants. However the restraint conferred by spared synaptic branches is still active in the absence of Hiw, since Wnd signaling can be further elevated by axotomy of all synapses in *hiw* null mutants (**Figure 3**). In a previous study we noted that the nerve crush injury led to a rapid down-regulation of an ectopically expressed Hiw-GFP transgene, and speculated that impairment of restraint by Hiw leads to activation of Wnd signaling in injured axons (Xiong et al., 2010). Our current data do not rule out a role for Hiw, but suggest the existence of additional mechanisms that restrain Wnd signaling at intact synapses. This has also been suggested from developmental studies of photoreceptor growth cone termination, in which Hiw-independent downregulation of Wnd protein occurs concomitantly with the development of presynaptic boutons (Feoktistov and Herman, 2016).

We speculate that the regulation of Wnd is linked to the trafficking of organelles between the synaptic terminal and cell body, akin to neurotrophin signaling, which relies on retrograde trafficking of signaling endosomes in axons (Cosker et al., 2008). Consistent with this idea, DLK signaling becomes activated following nerve growth factor withdrawal from distal axons (Ghosh et al., 2011; Larhammar et al., 2017). Previous studies of Wnd signaling have documented its dependence on retrograde axonal transport machinery (Xiong et al., 2010). Moreover, mutations that disrupt axonal cytoskeleton and the unc-104/kif1A kinesin also lead to Wnd/DLK signaling activation (Li et al., 2017; Valakh et al., 2015, 2013). Both Wnd and its homologue DLK in mice show regulated association with organelle membranes via palmitoylation, and disruption of its palmitoylation abolishes DLK’s signaling ability (Holland et al., 2016; Kim et al., 2024; Niu et al., 2022). Palmitoylation and depalmitoylation are dynamically regulated in axons (Ramzan et al., 2023; Zhang et al., 2024), hence comprise an attractive mechanism for mediating restraint of DLK at synaptic branches. Future delineation of the organelle(s) that Wnd/DLK associates with may provide important clues to its mechanism of regulation.

Our observations that Wnd signaling could be restrained by an intact axonal bifurcation suggests that at least one level of regulation could occur in the cell body. Consistent with this idea, a recent study has shown that Wnd signaling can be ectopically activated in the cell body when its transport to distal synapses is impaired in *rab11* mutants(Kim et al., 2024). Given the many factors that have been documented thus far that regulate DLK/Wnd protein or signaling(Asghari Adib et al., 2018), we think that this kinase must be tightly regulated both at synapses and cell bodies (**Figure 4**). Regulation at both locations gives the neuron a way to monitor the state of its entire axon and restrict signaling activation to scenarios where all efferent connections of the axon are disrupted.

**Figure 4:**
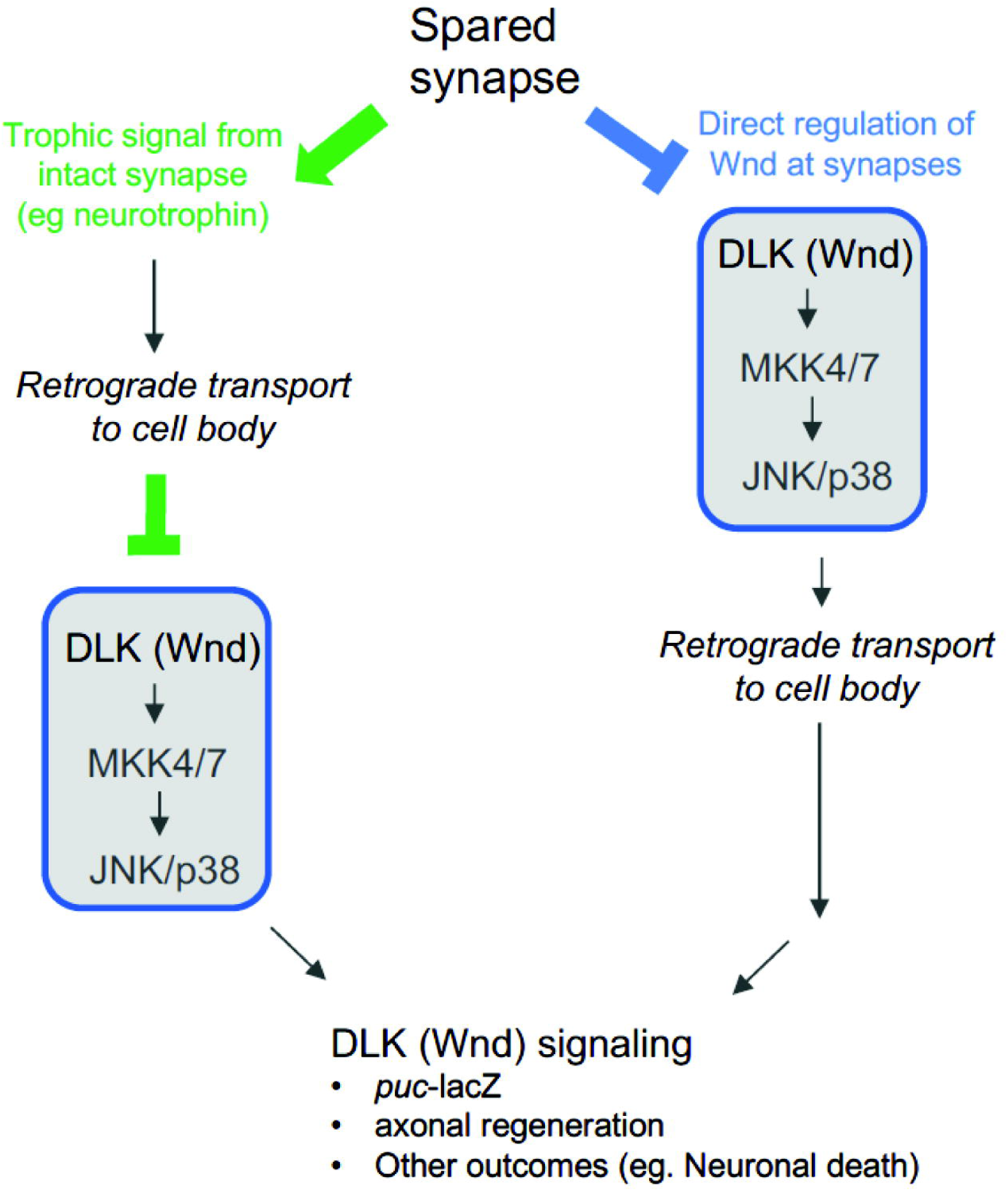
Potential mechanisms for regulation of Wnd signaling from synaptic terminals. In green, Wnd signaling may be regulated in the cell body downstream of a retrogradely transported signal. (For example, neurotrophin signaling). In blue, Wnd signaling activation is restrained locally at synaptic terminals, perhaps by regulating the levels or activation of Wnd itself. Activated Wnd or a downstream signaling factor is then retrogradely transported to the cell body. Previous observations that inhibition of retrograde transport blocks the induction of Wnd signaling following axonal injury favors the latter (blue) possibility. However the restraint conferred by a physically separate bifurcation suggests that an inhibitor of Wnd signaling activation can be retrogradely transported (green). We speculate that both mechanisms act as dual checkpoints to restrain Wnd signaling activation in the context of healthy circuits.

## Materials and Methods

### Drosophila stocks/Genetics

All fly crosses were raised at 25°C in a 12h light/dark cycle on standard sucrose and yeast food. Both male and female larvae were used unless otherwise stated. The following strains were used in this study: BL labeled flies were obtained from Bloomington Drosophila Stock Center (Indiana University), v labeled flies were obtained from the Vienna Drosophila Resource Center (VDRC, Vienna Biocenter Core Facilities). UAS-mCD8-GFP(Lee and Luo, 1999) (RRID:BDSC_5137), m12-gal4 (P(Gal4)^5053A^)(Ritzenthaler et al., 2000), (RRID:BDSC_2702), BG380-gal4 (C80-Gal4)(Budnik et al., 1996; Sanyal, 2009), puc-lacZ^E69^ (Martín-Blanco et al., 1998), *hiw*^Δ*N*^ (Wu et al., 2005)(RRID:BDSC_51637). UAS-Bitbow2 (Li et al., 2021), Tdc2-Gal4 (RRID:BDSC_9313), UAS-*wnd* RNAi (RRID:BDSC_35369), UAS-LexA RNAi (RRID:BDSC_67947). Additional stocks tested are listed in **Supplementary Table 1**.

### Immunohistochemistry

Wandering second and third instar larvae were dissected in ice-cold 1x PBS, fixed in 4% paraformaldehyde (16% diluted in 1X PBS, Electron microscopy Labs) for 20 mins at room temperature and washed thrice in 1X PBS. Tissues were blocked for a minimum 30 mins in 5% normal goat serum (NGS) in 0.25% triton X 100 in 1X PBS (PBST). Primary antibodies were incubated overnight in 5% NGS in 0.25% PBST at room temperature unless otherwise stated. The following antibodies and concentrations were used: mouse anti-lacZ (1:100, DSHB Cat# 40-1a, RRID:AB_528100), rabbit anti-DsRed (1:1000, Takara Bio Cat# 632496, RRID:AB_10013483), A488 rabbit anti-GFP (1:1000, for 2 hours at room temperature, Molecular Probes Cat# A-21311, RRID:AB_221477). Tissues were washed thrice in 1X PBS and incubated at room temperature for 2 hours in the appropriate secondary antibodies (1:1000 in 5% NGS in 0.25% PBST): AlexFluor 568 goat anti-mouse (ThermoFisher, A11004, RRID:AB_2534072), AlexFluor 488 goat anti-mouse, (ThermoFisher A32723, RRID:AB_2633275), AlexFluor 568 goat anti-rabbit, ThermoFisher A-11011, RRID:AB_143157). Tissues were washed thrice in 1x PBS and mounted onto superfrost plus slides (Fisher Scientific) and coverslipped with ProLong Diamond Antifade media (ThermoFisher Scientific, P36970). All experimental and control larval groups were processed (dissected, fixed, stained and images captured) together using identical confocal settings.

### Nerve crush assays

Peripheral nerve crushes in wandering 2nd instar larvae were performed as previously described(Waller et al., 2024; Xiong et al., 2010). Briefly, with the ventral side facing upwards, anesthetized larvae (taken from 1X PBS and placed on CO_2_ pad for ∼3-5 mins) had a small region of the cuticle (around abdominal segment 2) with underlying segmental nerves gently pinched with a pair of Dumont #5 fine forceps (Roboz Surgical RS4978). After injury, larvae were placed in small dishes containing standard fly food and kept at 25°C (12h light/dark) for 24h. Injuries were confirmed by the presence of posterior tail paralysis. For the new spared nerve crush assay, we performed injuries on wandering 2nd instar larvae with labeled type II octopaminergic neurons driven by Tdc2-gal4. The Tdc2-gal4 drives expression in 3 ventrally located midline neurons that sends one axon to the left side and one axon to the right side of the larvae. Similarly to complete nerve crushes, anesthetized larvae (ventral side up) were gently pinched with fine forceps on the left side (around abdominal segment 2), injuring nerves on one side while sparing nerves on the other side. Animals were placed in small dishes containing standard fly food and kept at 25°C (12h light/dark) for 24-72 hours.

### Axon/synaptic branch laser injury

As previously described (Ghannad-Rezaie et al., 2012; Mishra et al., 2014; Smithson et al., 2024b; Waller et al., 2024), single wandering 2^nd^ instar larva were taken from a dish containing 1X PBS and gently placed onto a kim wipe to remove excess PBS and then dipped into halocarbon oil 700 (H8898, Sigma). Each larva was then placed onto a glass coverslip dorsal side up (for inverted confocal) with an anterior to posterior (left to right) orientation. A PDMS microfluidic chip (https://www.ukosmos.com/) was placed on top of the larvae applying light force. A tight seal was created by applying gentle suction from a 30-ml syringe plunger. This vacuum suction immobilizes the intact larvae. The coverslip containing the microfluidic chip and larvae was mounted onto an Improvision spinning disk confocal system (PerkinElmer) connected to a Micropoint Laser Illumination and Ablation System (Andor Technology). The method for laser-induced microsurgery is described in (Smithson et al., 2024a). Prior to injury, the laser strength and region of interest were calibrated and optimized. Individual axonal branches (membrane GFP labeled) were identified and laser ablated. Confirmation of injury was demonstrated by a small absence of membrane GFP and degeneration of the proximal axon was detected as early as 8 hours after laser injury.

### Imaging, Quantification and Analysis

Both experimental and control larvae were imaged on an Improvision spinning disk confocal system (PerkinElmer) and quantified using identical confocal settings. To quantify *puc*-lacZ levels, the mean intensity of lacZ immunolabeling was measured in specific injury confirmed neurons. Injured m12-gal4 and Tdc2-gal4 axons were traced back to identify corresponding cell bodies in the lateral and medial ventral nerve cord. *Puc*-lacZ contains a NLS sequence fused to lacZ, resulting in nuclear expression of puc. The mean lacZ intensity was measured per neuron and normalized to uninjured control neurons. All imaging, image contrast/color adjustments, and quantification were conducted using Volocity software 6.2 (Improvision, Perkin Elmer). Data were analyzed and statistical tests were performed using Prism (GraphPad) and presented as mean ± SD.

## Supporting information

Merged supplemental materials

## Acknowledgements

We would like to thank Mariana Jimenez, Luis Rivera, and Sarah Cooke for help with tissue processing, Eric Robertson and Chris Jasinski for technical assistance. We thank Heather Broihier, Dion Dickman, Dhananjay Yellajoshyula and Jerry Silver for helpful discussions and comments on the manuscript. We thank Yuanquan Song (University of Pennsylvania) for sharing *piezo* flies, Yujia (Henry) Hu and Bing Ye (University of Michigan) for gtACR1 flies and technical advice. Stocks obtained from the Bloomington Drosophila Stock Center (NIH P40OD018537) and Vienna Drosophila Resource Center (VDRC, www.vdrc.at (Dietzl et al., 2007) were used in this study. This research was funded by the National Institutes for Health (NS069844 to CAC) and by the Canadian Institutes for Health Research (CIHR Fellowship to LJS).

## Notes

### Competing Interest Statement

The authors have declared no competing interest.

### Summary of Updates

The revision shows the branched injury data in Figure 1 in more extended form to enable better comparison of branched injuries at different locations, which remove varying numbers of synaptic boutons.

## References Cited

Abrams B, Grill B, Huang X, Jin Y. 2008. Cellular and molecular determinants targeting the Caenorhabditis elegans PHR protein RPM-1 to perisynaptic regions. Dev Dyn 237:630– 639.

Asghari Adib E, Smithson LJ, Collins CA. 2018. An axonal stress response pathway: degenerative and regenerative signaling by DLK. Curr Opin Neurobiol 53:110–119.

Bhattacharya MRC, Gerdts J, Naylor SA, Royse EX, Ebstein SY, Sasaki Y, Milbrandt J, DiAntonio A. 2012. A model of toxic neuropathy in Drosophila reveals a role for MORN4 in promoting axonal degeneration. J Neurosci 32:5054–5061.

Bounoutas A, Kratz J, Emtage L, Ma C, Nguyen KC, Chalfie M. 2011. Microtubule depolymerization in *Caenorhabditis elegans* touch receptor neurons reduces gene expression through a p38 MAPK pathway. Proc Natl Acad Sci U S A 108:3982–3987.

Budnik V, Koh YH, Guan B, Hartmann B, Hough C, Woods D, Gorczyca M. 1996. Regulation of synapse structure and function by the Drosophila tumor suppressor gene dlg. Neuron 17:627–640.

Chen C-H, Lee A, Liao C-P, Liu Y-W, Pan C-L. 2014. RHGF-1/PDZ-RhoGEF and retrograde DLK-1 signaling drive neuronal remodeling on microtubule disassembly. Proc Natl Acad Sci U S A 111:16568–16573.

Chen L, Wang Z, Ghosh-Roy A, Hubert T, Yan D, O’Rourke S, Bowerman B, Wu Z, Jin Y, Chisholm AD. 2011. Axon regeneration pathways identified by systematic genetic screening in C. elegans. Neuron 71:1043–1057.

Choi BJ, Imlach WL, Jiao W, Wolfram V, Wu Y, Grbic M, Cela C, Baines RA, Nitabach MN, McCabe BD. 2014. Miniature neurotransmission regulates Drosophila synaptic structural maturation. Neuron 82:618–634.

Collins CA, Wairkar YP, Johnson SL, DiAntonio A. 2006. Highwire restrains synaptic growth by attenuating a MAP kinase signal. Neuron 51:57–69.

Cosker KE, Courchesne SL, Segal RA. 2008. Action in the axon: generation and transport of signaling endosomes. Curr Opin Neurobiol 18:270–275.

Daniels RW, Collins CA, Chen K, Gelfand MV, Featherstone DE, DiAntonio A. 2006. A single vesicular glutamate transporter is sufficient to fill a synaptic vesicle. Neuron 49:11–16.

DeVault L, Mateusiak C, Palucki J, Brent M, Milbrandt J, DiAntonio A. 2024. The response of Dual-leucine zipper kinase (DLK) to nocodazole: Evidence for a homeostatic cytoskeletal repair mechanism. PLoS One 19:e0300539.

DiAntonio A, Schwarz TL. 1994. The effect on synaptic physiology of synaptotagmin mutations in Drosophila. Neuron 12:909–920.

Dietzl G, Chen D, Schnorrer F, Su K-C, Barinova Y, Fellner M, Gasser B, Kinsey K, Oppel S, Scheiblauer S, Couto A, Marra V, Keleman K, Dickson BJ. 2007. A genome-wide transgenic RNAi library for conditional gene inactivation in Drosophila. Nature 448:151– 156.

Feoktistov AI, Herman TG. 2016. Wallenda/DLK protein levels are temporally downregulated by Tramtrack69 to allow R7 growth cones to become stationary boutons. Development 143:2983–2993.

Fernandes KJ, Fan DP, Tsui BJ, Cassar SL, Tetzlaff W. 1999. Influence of the axotomy to cell body distance in rat rubrospinal and spinal motoneurons: differential regulation of GAP-43, tubulins, and neurofilament-M. J Comp Neurol 414:495–510.

Ghannad-Rezaie M, Wang X, Mishra B, Collins C, Chronis N. 2012. Microfluidic chips for in vivo imaging of cellular responses to neural injury in Drosophila larvae. PLoS One 7:e29869.

Ghosh AS, Wang B, Pozniak CD, Chen M, Watts RJ, Lewcock JW. 2011. DLK induces developmental neuronal degeneration via selective regulation of proapoptotic JNK activity. J Cell Biol 194:751–764.

Guedes-Dias P, Holzbaur ELF. 2019. Axonal transport: Driving synaptic function. Science 366. doi:10.1126/science.aaw9997

Hammarlund M, Nix P, Hauth L, Jorgensen EM, Bastiani M. 2009. Axon regeneration requires a conserved MAP kinase pathway. Science 323:802–806.

Han Y, Chien C, Goel P, He K, Pinales C, Buser C, Dickman D. 2022. Botulinum neurotoxin accurately separates tonic vs. phasic transmission and reveals heterosynaptic plasticity rules in Drosophila. Elife 11. doi:10.7554/eLife.77924

Hilton BJ, Husch A, Schaffran B, Lin T-C, Burnside ER, Dupraz S, Schelski M, Kim J, Müller JA, Schoch S, Imig C, Brose N, Bradke F. 2022. An active vesicle priming machinery suppresses axon regeneration upon adult CNS injury. Neuron 110:51–69.e7.

Holland SM, Collura KM, Ketschek A, Noma K, Ferguson TA, Jin Y, Gallo G, Thomas GM. 2016. Palmitoylation controls DLK localization, interactions and activity to ensure effective axonal injury signaling. Proc Natl Acad Sci U S A 113:763–768.

Huebner EA, Strittmatter SM. 2009. Axon regeneration in the peripheral and central nervous systems. Results Probl Cell Differ 48:339–351.

Huntwork-Rodriguez S, Wang B, Watkins T, Ghosh AS, Pozniak CD, Bustos D, Newton K, Kirkpatrick DS, Lewcock JW. 2013. JNK-mediated phosphorylation of DLK suppresses its ubiquitination to promote neuronal apoptosis. J Cell Biol 202:747–763.

Hu Z, Deng N, Liu K, Zeng W. 2019. DLK mediates the neuronal intrinsic immune response and regulates glial reaction and neuropathic pain. Exp Neurol 322:113056.

Joy MT, Ben Assayag E, Shabashov-Stone D, Liraz-Zaltsman S, Mazzitelli J, Arenas M, Abduljawad N, Kliper E, Korczyn AD, Thareja NS, Kesner EL, Zhou M, Huang S, Silva TK, Katz N, Bornstein NM, Silva AJ, Shohami E, Carmichael ST. 2019. CCR5 Is a Therapeutic Target for Recovery after Stroke and Traumatic Brain Injury. Cell 176:1143–1157.e13.

Kim MD, Wen Y, Jan Y-N. 2009. Patterning and organization of motor neuron dendrites in the Drosophila larva. Dev Biol 336:213–221.

Kim SE, Coste B, Chadha A, Cook B, Patapoutian A. 2012. The role of Drosophila Piezo in mechanical nociception. Nature 483:209–212.

Kim S, Quagraine Y, Singh M, Kim JH. 2024. Rab11 suppresses neuronal stress signaling by localizing Dual leucine zipper kinase to axon terminals for protein turnover. bioRxiv. doi:10.1101/2023.04.18.537392

Kitamoto T. 2001. Conditional modification of behavior in Drosophila by targeted expression of a temperature-sensitive shibire allele in defined neurons. J Neurobiol 47:81–92.

Koon AC, Ashley J, Barria R, DasGupta S, Brain R, Waddell S, Alkema MJ, Budnik V. 2011. Autoregulatory and paracrine control of synaptic and behavioral plasticity by octopaminergic signaling. Nat Neurosci 14:190–199.

Kurup N, Yan D, Goncharov A, Jin Y. 2015. Dynamic microtubules drive circuit rewiring in the absence of neurite remodeling. Curr Biol 25:1594–1605.

Larhammar M, Huntwork-Rodriguez S, Rudhard Y, Sengupta-Ghosh A, Lewcock JW. 2017. The Ste20 Family Kinases MAP4K4, MINK1, and TNIK Converge to Regulate Stress-Induced JNK Signaling in Neurons. J Neurosci 37:11074–11084.

Lee T, Luo L. 1999. Mosaic analysis with a repressible cell marker for studies of gene function in neuronal morphogenesis. Neuron 22:451–461.

Le Pichon CE, Meilandt WJ, Dominguez S, Solanoy H, Lin H, Ngu H, Gogineni A, Sengupta Ghosh A, Jiang Z, Lee S-H, Maloney J, Gandham VD, Pozniak CD, Wang B, Lee S, Siu M, Patel S, Modrusan Z, Liu X, Rudhard Y, Baca M, Gustafson A, Kaminker J, Carano RAD, Huang EJ, Foreman O, Weimer R, Scearce-Levie K, Lewcock JW. 2017. Loss of dual leucine zipper kinase signaling is protective in animal models of neurodegenerative disease. Sci Transl Med 9. doi:10.1126/scitranslmed.aag0394

Li J, Zhang YV, Asghari Adib E, Stanchev DT, Xiong X, Klinedinst S, Soppina P, Jahn TR, Hume RI, Rasse TM, Collins CA. 2017. Restraint of presynaptic protein levels by Wnd/DLK signaling mediates synaptic defects associated with the kinesin-3 motor Unc-104. Elife 6. doi:10.7554/eLife.24271

Li Y, Walker LA, Zhao Y, Edwards EM, Michki NS, Cheng HPJ, Ghazzi M, Chen TY, Chen M, Roossien DH, Cai D. 2021. Bitbow Enables Highly Efficient Neuronal Lineage Tracing and Morphology Reconstruction in Single Drosophila Brains. Front Neural Circuits 15:732183.

Lorenzana AO, Lee JK, Mui M, Chang A, Zheng B. 2015. A surviving intact branch stabilizes remaining axon architecture after injury as revealed by in vivo imaging in the mouse spinal cord. Neuron 86:947–954.

Martín-Blanco E, Gampel A, Ring J, Virdee K, Kirov N, Tolkovsky AM, Martinez-Arias A. 1998. puckered encodes a phosphatase that mediates a feedback loop regulating JNK activity during dorsal closure in Drosophila. Genes Dev 12:557–570.

Mason MRJ, Lieberman AR, Anderson PN. 2003. Corticospinal neurons up-regulate a range of growth-associated genes following intracortical, but not spinal, axotomy. Eur J Neurosci 18:789–802.

Mishra B, Ghannad-Rezaie M, Li J, Wang X, Hao Y, Ye B, Chronis N, Collins CA. 2014. Using microfluidics chips for live imaging and study of injury responses in Drosophila larvae. J Vis Exp e50998.

Mohammad F, Stewart JC, Ott S, Chlebikova K, Chua JY, Koh T-W, Ho J, Claridge-Chang A. 2017. Optogenetic inhibition of behavior with anion channelrhodopsins. Nat Methods 14:271–274.

Monastirioti M, Gorczyca M, Rapus J, Eckert M, White K, Budnik V. 1995. Octopamine immunoreactivity in the fruit fly Drosophila melanogaster. J Comp Neurol 356:275–287.

Mosca TJ, Carrillo RA, White BH, Keshishian H. 2005. Dissection of synaptic excitability phenotypes by using a dominant-negative Shaker K+ channel subunit. Proc Natl Acad Sci U S A 102:3477–3482.

Nakata K, Abrams B, Grill B, Goncharov A, Huang X, Chisholm AD, Jin Y. 2005. Regulation of a DLK-1 and p38 MAP kinase pathway by the ubiquitin ligase RPM-1 is required for presynaptic development. Cell 120:407–420.

Nitabach MN, Blau J, Holmes TC. 2002. Electrical silencing of Drosophila pacemaker neurons stops the free-running circadian clock. Cell 109:485–495.

Niu J, Holland SM, Ketschek A, Collura KM, Hesketh NL, Hayashi T, Gallo G, Thomas GM. 2022. Palmitoylation couples the kinases DLK and JNK3 to facilitate prodegenerative axon-to-soma signaling. Sci Signal 15:eabh2674.

Opperman KJ, Grill B. 2014. RPM-1 is localized to distinct subcellular compartments and regulates axon length in GABAergic motor neurons. Neural Dev 9:10.

Patel S, Meilandt WJ, Erickson RI, Chen J, Deshmukh G, Estrada AA, Fuji RN, Gibbons P, Gustafson A, Harris SF, Imperio J, Liu W, Liu X, Liu Y, Lyssikatos JP, Ma C, Yin J, Lewcock JW, Siu M. 2017. Selective Inhibitors of Dual Leucine Zipper Kinase (DLK, MAP3K12) with Activity in a Model of Alzheimer’s Disease. J Med Chem 60:8083–8102.

Pérez-Moreno JJ, O’Kane CJ. 2019. GAL4 Drivers Specific for Type Ib and Type Is Motor Neurons in Drosophila. G3 9:453–462.

Petzoldt AG. 2023. Presynaptic Precursor Vesicles-Cargo, Biogenesis, and Kinesin-Based Transport across Species. Cells 12. doi:10.3390/cells12182248

Pittendrigh B, Reenan R, ffrench-Constant RH, Ganetzky B. 1997. Point mutations in the Drosophila sodium channel gene para associated with resistance to DDT and pyrethroid insecticides. Mol Gen Genet 256:602–610.

Raineteau O, Schwab ME. 2001. Plasticity of motor systems after incomplete spinal cord injury. Nat Rev Neurosci 2:263–273.

Ramón y Cajal S. 1928. Degeneration & Regeneration of the Nervous System, By S. Ramon Y Cajal. Translated and Edited by Raoul M. May.

Ramzan F, Abrar F, Mishra GG, Liao LMQ, Martin DDO. 2023. Lost in traffic: consequences of altered palmitoylation in neurodegeneration. Front Physiol 14:1166125.

Ritzenthaler S, Suzuki E, Chiba A. 2000. Postsynaptic filopodia in muscle cells interact with innervating motoneuron axons. Nat Neurosci 3:1012–1017.

Sanyal S. 2009. Genomic mapping and expression patterns of C380, OK6 and D42 enhancer trap lines in the larval nervous system of Drosophila. *Gene* Expr Patterns 9:371–380.

Smithson LJ, Waller TJ, Collins CA. 2024a. Laser Microsurgery in Drosophila Larvae Using the MicroPoint Ablation System. Cold Spring Harb Protoc. doi:10.1101/pdb.prot108171

Smithson LJ, Waller TJ, Collins CA. 2024b. Immobilizing Second-Instar Drosophila Larvae for Imaging and Surgery Using the Larva Chip. Cold Spring Harb Protoc. doi:10.1101/pdb.prot108170

Stone MC, Albertson RM, Chen L, Rolls MM. 2014. Dendrite injury triggers DLK-independent regeneration. Cell Rep 6:247–253.

Tedeschi A, Dupraz S, Laskowski CJ, Xue J, Ulas T, Beyer M, Schultze JL, Bradke F. 2016. The Calcium Channel Subunit Alpha2delta2 Suppresses Axon Regeneration in the Adult CNS. Neuron 92:419–434.

Valakh V, Frey E, Babetto E, Walker LJ, DiAntonio A. 2015. Cytoskeletal disruption activates the DLK/JNK pathway, which promotes axonal regeneration and mimics a preconditioning injury. Neurobiol Dis 77:13–25.

Valakh V, Walker LJ, Skeath JB, DiAntonio A. 2013. Loss of the spectraplakin short stop activates the DLK injury response pathway in Drosophila. J Neurosci 33:17863–17873.

Vömel M, Wegener C. 2008. Neuroarchitecture of aminergic systems in the larval ventral ganglion of Drosophila melanogaster. PLoS One 3:e1848.

Waller TJ, Smithson LJ, Collins CA. 2024. Peripheral Nerve Crush in Drosophila Larvae. Cold Spring Harb Protoc. doi:10.1101/pdb.prot108169

Wang Z, Kumaran M, Batsel E, Testor-Cabrera S, Beine Z, Ribelles AA, Tsoulfas P, Venkatesh I, Blackmore MG. 2024. Injury distance limits the transcriptional response to spinal injury. bioRxiv. doi:10.1101/2024.05.27.596075

Wan HI, DiAntonio A, Fetter RD, Bergstrom K, Strauss R, Goodman CS. 2000. Highwire regulates synaptic growth in Drosophila. Neuron 26:313–329.

Watkins TA, Wang B, Huntwork-Rodriguez S, Yang J, Jiang Z, Eastham-Anderson J, Modrusan Z, Kaminker JS, Tessier-Lavigne M, Lewcock JW. 2013. DLK initiates a transcriptional program that couples apoptotic and regenerative responses to axonal injury. Proc Natl Acad Sci U S A 110:4039–4044.

Welsbie DS, Mitchell KL, Jaskula-Ranga V, Sluch VM, Yang Z, Kim J, Buehler E, Patel A, Martin SE, Zhang P-W, Ge Y, Duan Y, Fuller J, Kim B-J, Hamed E, Chamling X, Lei L, Fraser IDC, Ronai ZA, Berlinicke CA, Zack DJ. 2017. Enhanced Functional Genomic Screening Identifies Novel Mediators of Dual Leucine Zipper Kinase-Dependent Injury Signaling in Neurons. Neuron 94:1142–1154.e6.

Welsbie DS, Yang Z, Ge Y, Mitchell KL, Zhou X, Martin SE, Berlinicke CA, Hackler L Jr, Fuller J, Fu J, Cao LH, Han B, Auld D, Xue T, Hirai S, Germain L, Simard-Bisson C, Blouin R, Nguyen JV, Davis CH, Enke RA, Boye SL, Merbs SL, Marsh-Armstrong N, Hauswirth WW, DiAntonio A, Nickells RW, Inglese J, Hanes J, Yau KW, Quigley HA, Zack DJ. 2013. Functional genomic screening identifies dual leucine zipper kinase as a key mediator of retinal ganglion cell death. Proc Natl Acad Sci U S A 110:4045–4050.

Wlaschin JJ, Gluski JM, Nguyen E, Silberberg H, Thompson JH, Chesler AT, Le Pichon CE. 2018. Dual leucine zipper kinase is required for mechanical allodynia and microgliosis after nerve injury. Elife 7. doi:10.7554/eLife.33910

Wu C, Wairkar YP, Collins CA, DiAntonio A. 2005. Highwire function at the Drosophila neuromuscular junction: spatial, structural, and temporal requirements. J Neurosci 25:9557–9566.

Wu Z, Ghosh-Roy A, Yanik MF, Zhang JZ, Jin Y, Chisholm AD. 2007. *Caenorhabditis elegans* neuronal regeneration is influenced by life stage, ephrin signaling, and synaptic branching. Proc Natl Acad Sci U S A 104:15132–15137.

Xiong X, Collins CA. 2012. A conditioning lesion protects axons from degeneration via the Wallenda/DLK MAP kinase signaling cascade. J Neurosci 32:610–615.

Xiong X, Hao Y, Sun K, Li J, Li X, Mishra B, Soppina P, Wu C, Hume RI, Collins CA. 2012. The Highwire ubiquitin ligase promotes axonal degeneration by tuning levels of Nmnat protein. PLoS Biol 10:e1001440.

Xiong X, Wang X, Ewanek R, Bhat P, Diantonio A, Collins CA. 2010. Protein turnover of the Wallenda/DLK kinase regulates a retrograde response to axonal injury. J Cell Biol 191:211– 223.

Yan D, Wu Z, Chisholm AD, Jin Y. 2009. The DLK-1 kinase promotes mRNA stability and local translation in C. elegans synapses and axon regeneration. Cell 138:1005–1018.

Zhang X, Jeong H, Niu J, Holland SM, Rotanz BN, Gordon J, Einarson MB, Childers WE, Thomas GM. 2024. Novel inhibitors of acute, axonal DLK palmitoylation are neuroprotective and avoid the deleterious side effects of cell-wide DLK inhibition. bioRxiv. doi:10.1101/2024.04.19.590310

Zhen M, Huang X, Bamber B, Jin Y. 2000. Regulation of presynaptic terminal organization by C. elegans RPM-1, a putative guanine nucleotide exchanger with a RING-H2 finger domain. Neuron 26:331–343.

