## Supplementary material for "Axonal injury signaling is restrained by a spared synaptic branch": Merged supplemental materials

**Supplemental Figure 1: Further characterization of branched injury assays to SNc and aCC motoneurons.**

**A)** cartoon denoting nomenclature of MNSNc-26/29 (red) and MNSNc-27 (blue) branches.

**B**) Example images of nerve terminals and cell bodies of m12-Gal4; UAS-Bitbow2 labeled MNSNc neurons, used to confirm the anatomy.

**C)** Similarly to laser axotomy in **Figure 1C**, nerve crush injury (24h) induces *puc*-lacZ expression in both MNSNc neurons, but not in MNSNc neurons that co-express *wnd*-RNAi.

**D)** Synaptic terminals (top row) and cell bodies (bottom row) of aCC motoneurons on muscle 1, labeled in Dpr4-Gal4, UAS-mCD8-GFP larvae. *puc*-lacZ expression (red) is induced following injuries that result in loss of all synaptic boutons but not following injuries to one branch that leave the other branch intact.

**E)** Quantification of *puc*-lacZ intensities in aCC neurons. A one-way ANOVA with Tukey test for multiple comparisons was performed. **** $p < 0.0001$; ns = not significant.

**F-G)** Full (F) but not partial (G) removal of synaptic branches induces stability and trafficking ectopically expressed kinase-dead GFP-Wnd-KD (in UAS-GFP-Wnd-KD; m12-Gal4, UAS-mCD8-RFP animals). Synaptic branches from muscle 27 (F), or muscle 29 (G) were axotomized by laser surgery and imaged following 24h.

**F)** GFP-Wnd-KD protein accumulates at the proximal tip of axons that have lost all synaptic boutons.

**G)** GFP-Wnd-KD is barely detectable in axons following injuries that leave spared synaptic branches. Not shown, GFP-Wnd-KD levels in G are similar in uninjured MNSNc axons.

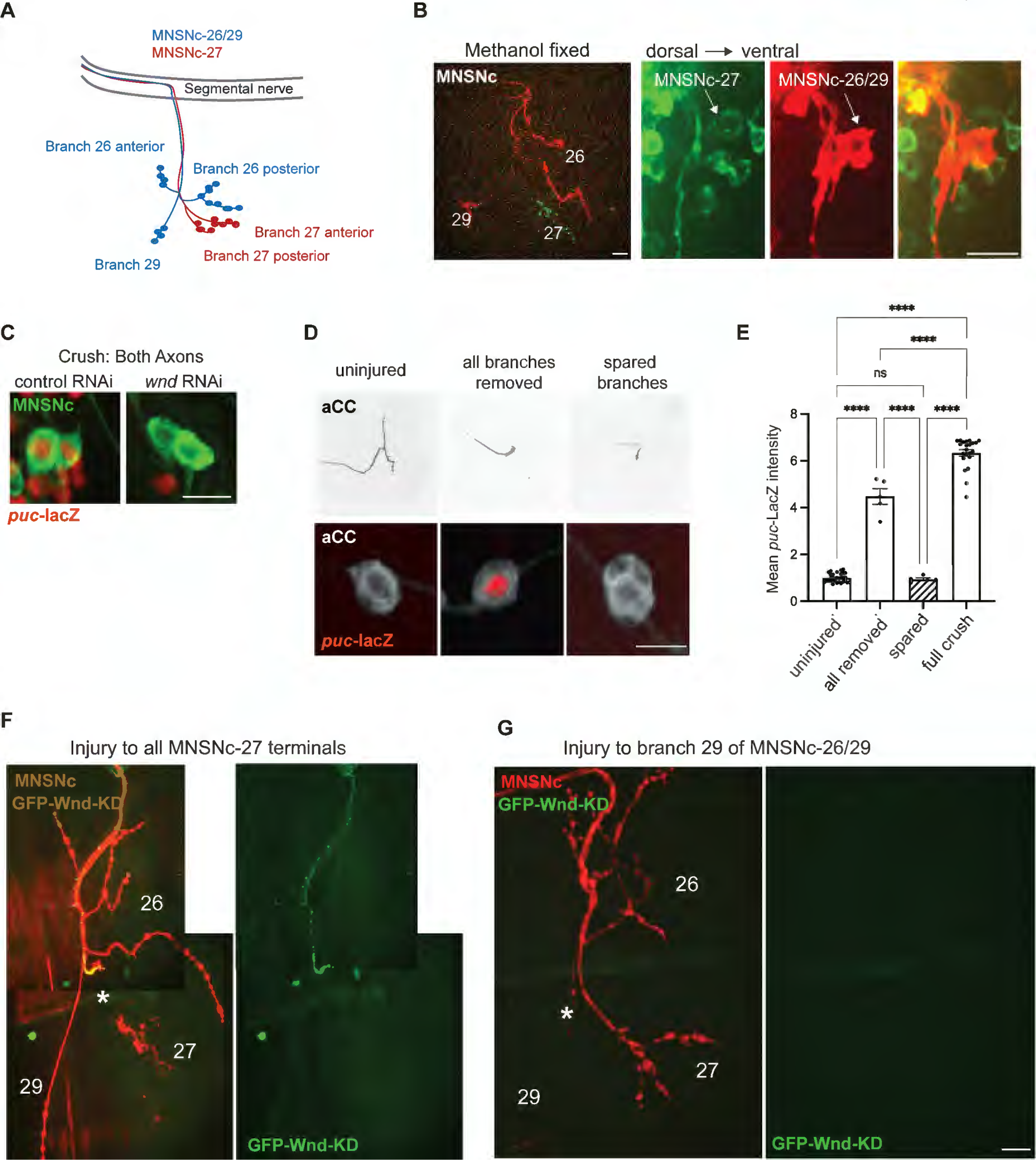
A
MNSNc-26/29
MNSNc-27
Segmental nerve
Branch 26 anterior
Branch 26 posterior
Branch 27 anterior
Branch 27 posterior
Branch 29
B
Methanol fixed
dorsal → ventral
MNSNc
26
29
27
MNSNc-27
MNSNc-26/29
C
Crush: Both Axons
control RNAi
wnd RNAi
MNSNc
puc-lacZ
D
uninjured
all branches removed
spared branches
aCC
aCC
puc-lacZ
E
Mean puc-LacZ intensity
0
2
4
6
8
****
****
ns
****
****
****
uninjured
all removed
spared
full crush
F
Injury to all MNSNc-27 terminals
MNSNc
GFP-Wnd-KD
26
*
27
29
GFP-Wnd-KD
G
Injury to branch 29 of MNSNc-26/29
MNSNc
GFP-Wnd-KD
26
*
27
29
GFP-Wnd-KD

**Supplemental Figure 2: Anatomy and laser surgery of Tdc2 bifurcated neurons.**

**A-E)** Views of Tdc2-Gal4, UAS-mCD8-GFP expressing VUM neurons to illustrate their anatomy.

**A)** Individual confocal plane that shows 3 cell bodies in individual segments, which lie in the middle of the nerve cord. **B)** Side view and **C)** top view that show the locations of the bifurcations.

**D)** Cartoon of the 3 neurons from one segment, showing their bifurcations to symmetrical sides of the animal.

**E)** composite and *camera lucida* views of the NMJ terminals for the 3 neurons on one abdominal hemisegment. The 3 Tdc2 neurons each form stereotyped branches to innervate a unique group of muscles.

**F)** Example results from laser axotomies to individual Tdc2/VUM neurons (24h after injury) on either one side or both sides of the animal. Surgeries were carried out at the indicated locations which lie upstream of the final synaptic branches (at the transition zone between the segmental nerve and abdominal muscles). The stereotyped anatomy allows for identification of each VUM neuron (labeled 1,2 and 3). Injured branches are marked with an asterisk while spared are marked with squares. For each neuron, only injuries to both bifurcations allowed for induction of *puc*-lacZ. Scale bars = 20 um.

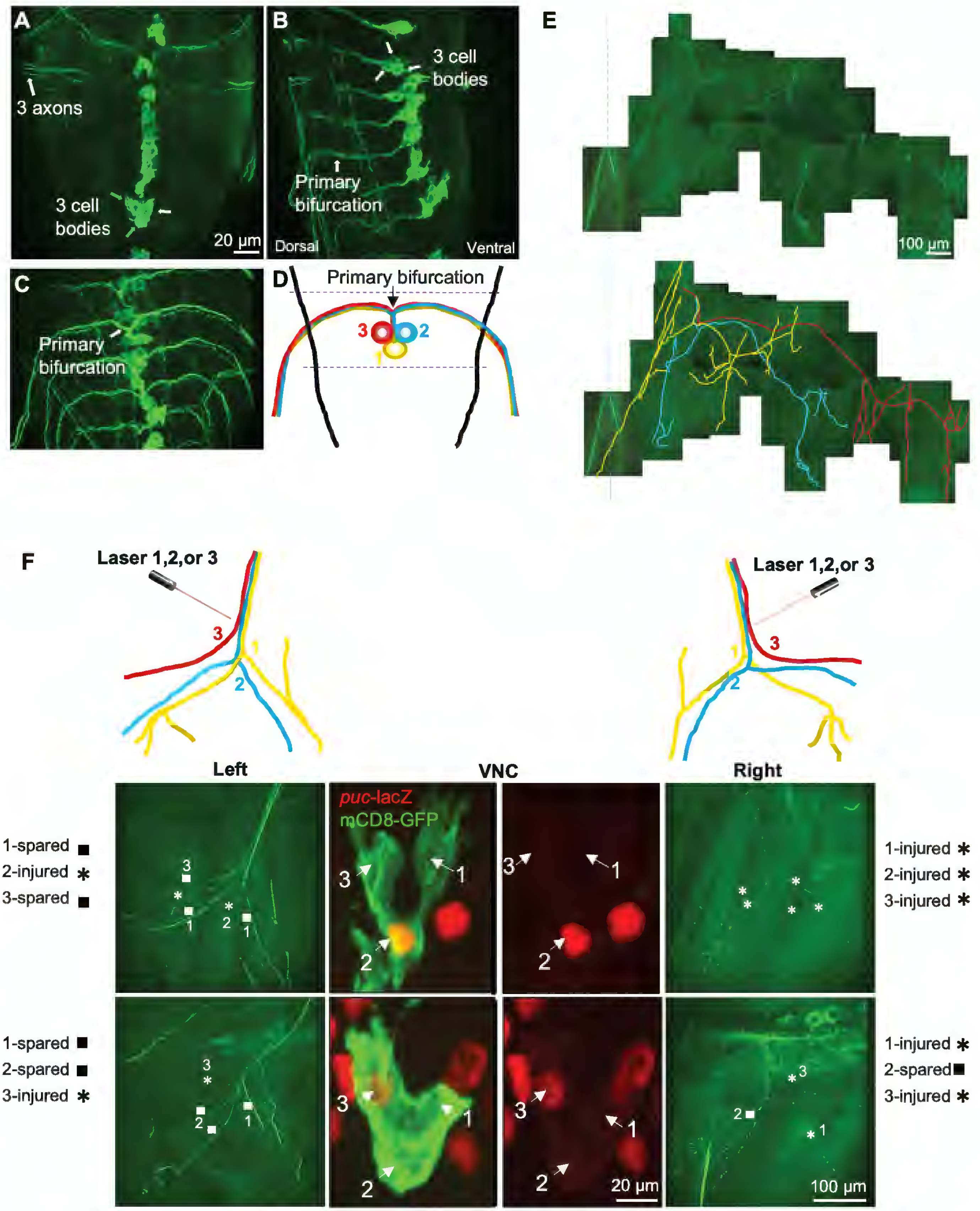
A
3 axons
3 cell bodies
20 μm
B
3 cell bodies
Primary bifurcation
Dorsal
Ventral
C
Primary bifurcation
D
Primary bifurcation
3
2
1
E
100 μm
F
Laser 1,2,or 3
3
2
1
Laser 1,2,or 3
3
2
1
Left
VNC
Right
puc-lacZ
mCD8-GFP
1-spared
2-injured
3-spared
1-injured
2-injured
3-injured
1-spared
2-spared
3-injured
1-injured
2-spared
3-injured
20 μm
100 μm

**Supplemental Figure 3: Confirmation that elevated *puc*-lacZ expression in *hiw* mutants requires Wnd.**

**A)** Top row shows *puc*-lacZ expression (red) together with a marker for nuclei (Draq5, gray). Control (lexA)-RNAi or wnd-RNAi UAS lines are driven by the BG380Gal4 driver in the background of control versus $hiw^{\Delta N}$. Bottom row shows example NMJs in these genotypes. Scale bars = 20 um.

**B)** Quantification of *puc*-lacZ expression in A. A one-way ANOVA with Tukey test for multiple comparisons was performed. **** $p < 0.0001$; ns = not significant.

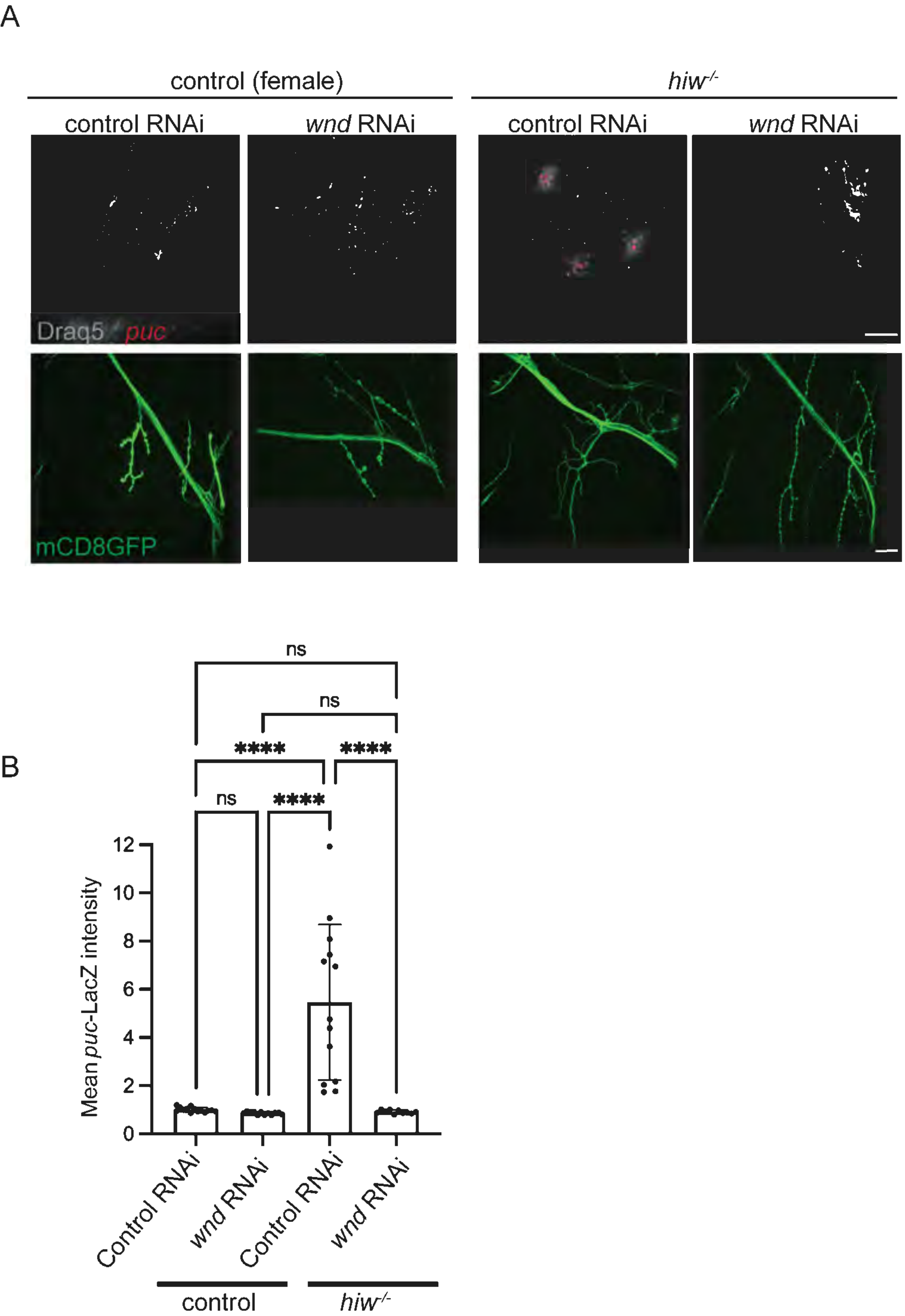
A
control (female)
hiw-/-
control RNAi
wnd RNAi
control RNAi
wnd RNAi
Draq5 puc
mCD8GFP
B
ns
ns
****
****
ns
****
Mean puc-LacZ intensity
0
2
4
6
8
10
12
Control RNAi
wnd RNAi
Control RNAi
wnd RNAi
control
hiw-/-

**Supplementary Table 1:** Summary of negative results from genetic manipulations that impair synaptic transmission and/or signaling at synapses. The listed UAS lines, UAS-RNAi lines and genetic mutations were tested for their ability to alter the expression of *puc*-lacZ in motoneurons. Outcomes and experimental details are noted.

| Name[LS1] | Stock ID | Target/Function | ref(s) | Driver (Gal4) | observation |
|---|---|---|---|---|---|
| **UAS lines** | | | | | |
| UAS-*shibire*-ts | RRID:BDSC_44222 | dominant temperature sensitive mutation in dynamin - confers rapid inhibition of synaptic vesicle endocytosis and exocytosis. | (Kitamoto, 2001) | m12 | temperature shift to 32C for 24h. (nSyb-Gal4, UAS-Shi-ts flies are paralyzed when shifted to 32C for 20 minutes). No difference between shifted (32C) and unshifted (25C) for uninjured, single branch injured, and complete axotomy. (shift was done immediately after laser surgery). |
| UAS-dORK-delta-C | RRID:BDSC_8928 | constitutively open K+ rectifying channel. Confers electrical silencing. | (Nitabach et al., 2002) | m12 | No difference from control (CS) flies, both uninjured and full axotomy. |
| UAS-dORK-delta-NC (non-conducting) | RRID:BDSC_6587 | Non-conducting version of UAS-ORK1 (control) | (Nitabach et al., 2002) | m12 | No difference from control (CS) flies, both uninjured and full axotomy. |
| UAS-gtACR1 | RRID:BDSC_92983 | light-inducible Cl- channel | (Mohammad et al., 2017) | m12 | Kept in dark until 24h before dissection, when larvae were shifted to 530 nm light (for 24h). No difference between vehicle (ethanol) and ATR (1mM) control. As a control for the method, BG380Gal4; UAS-gtACR1 (ATR) flies showed rapid paralysis with shifted 530 nm light, but animals raised with vehicle food did not. |
| UAS-*Sh*-DN | RRID:BDSC_40974 | mutated Shaker K+ channel which attenuates membrane excitability | (Mosca et al., 2005) | m12 | No difference from control (CS) flies, uninjured |

| | | | | | |
|---|---|---|---|---|---|
| UAS-tkv-CA | RRID:BDSC_36537 | consitutively activated type I BMP receptor (thickveins) | | BG380 | causes elevation of Gbb/BMP signaling (measured by pSMAD), but does not alter the induction of puc-lacZ in either injured (24h nerve crush) or uninjured animals. |
| **endogenous mutations** | | | | | |
| *paralytic (para)-ts* | | Temperature-sensitive allele of voltage-gated sodium channel | (Pittendrigh et al, 1997) | | tested 6h at 37C, and 24h at 29. No difference between shifted and unshifted for uninjured animals. Also no difference between shifted and unshifted for full axotomy. (shift was done immediately after nerve crush). |
| Dm-*piezo* KO | | *piezo* mechanosensitive Ca2+ channel | (Kim et al., 2012) | | Similar to control for *puc-lacZ* induction |
| **RNAi lines** | | | | | |
| *cacophony (cac)* | vdrc 101478 | alpha1 subunit of voltage-gated calcium channel | | m12 with dicer2 | no difference from UAS-control (luciferase) RNAi |
| | | | | BG380, Dicer2 | no difference from UAS-control (luciferase) RNAi |
| *straightjacket (stj)* | RRID:BDSC_25807 | alpha 2 delta subunit of voltage-gated calcium channel | | BG380, Dicer2 | no difference from UAS-control (luciferase) RNAi |
| | | | | D42 | no difference from UAS-control (luciferase) RNAi |
| *paralytic (para)* | RRID:BDSC_31471 | voltage-gated sodium channel | | BG380 | lethal |
| *paralytic (para)* | RRID:BDSC_31676 | voltage-gated sodium channel | | BG380, D42 | no difference from UAS-control (luciferase) RNAi |
| *shaker (sh)* | RRID:BDSC_31680 | shaker (sh) voltage gated K+ channel | | BG380, Dicer 2 | no difference from UAS-control (luciferase) RNAi |
| *shaker cognate (shaw)* | RRID:BDSC_28346 | voltage gated K channel Kv3.1 | | BG380, Dicer 2 | no difference from UAS-control (luciferase) RNAi |

| | | | | | |
|---|---|---|---|---|---|
| *synaptosomal-associated protein (SNAP25)* | RRID:BDSC_27306 | t-SNARE, regulation of neurotransmitter release | | BG380, Dicer2 | no difference from UAS-control (luciferase) RNAi |
| | RRID:BDSC_34377 | | | BG380, Dicer2 | no difference from UAS-control (luciferase) RNAi |
| | | | | D42 | no difference from UAS-control (luciferase) RNAi |
| *synaptobrevin (nSyb)* | RRID:BDSC_31983 | v-SNARE, role in synaptic vesicle exocytosis | | BG380, Dicer2 | no difference from UAS-control (luciferase) RNAi |
| | | | | m12 | no difference from UAS-control (luciferase) RNAi |
| *syntaxin 4 (syx4)* | RRID:BDSC_44054 | tSNARE, role in synaptic vesicle exocytosis | | BG380, Dicer2 | no difference from UAS-control (luciferase) RNAi |
| *syntaxin 1A (syx1A)* | RRID:BDSC_25811 | tSNARE, role in synaptic vesicle exocytosis | | BG380, Dicer2 | no difference from UAS-control (luciferase) RNAi |
| *synaptotagmin 4 (syt4)* | RRID:BDSC_39016 | calcium dependent exocytosis of secretory vesicles. | | BG380, Dicer2 | no difference from UAS-control (luciferase) RNAi |
| *vesicular glutamate transporter (vGlut)* | v104324 | vesicular glutamate transporter | | BG380, Dicer2 | no difference from UAS-control (luciferase) RNAi |
| | RRID:BDSC_40927 | | | D42 | no difference from UAS-control (luciferase) RNAi |
| | | | | BG380, Dicer2 | no difference from UAS-control (luciferase) RNAi |
| *piezo* | vdrc 2796 | mechanosensitive Ca2+ channel | | | no difference from UAS-control (luciferase) RNAi |
| *no mechanoreceptor potential C (nompC)* | vdrc 105579 | pore-forming subunit of mechanotransduction channel | | BG380, Dicer2 | no difference from UAS-control (luciferase) RNAi |
| *shortwing (sw)* | vdrc 101559 | intermediate chain subunit of dynein | | BG380, Dicer2 | no difference from UAS-control (luciferase) RNAi |
| | vdrc 48334 | | | BG380, Dicer2 | no difference from UAS-control (luciferase) RNAi |
| *neurexin 1 (nrx-1)* | vdrc 36326 | presynaptic protein, cell-cell interactions, exocytosis, synaptic signaling | | BG380, Dicer2 | no difference from UAS-control (luciferase) RNAi |
| | RRID:BDSC_27502 | | | BG380, Dicer2 | no difference from UAS-control (luciferase) RNAi |

| | | | | | |
|---|---|---|---|---|---|
| | | | | D42 | no difference from UAS-control (luciferase) RNAi |
| | RRID:BDSC_32408 | | | BG380, Dicer2 | no difference from UAS-control (luciferase) RNAi |
| | | | | D42 | no difference from UAS-control (luciferase) RNAi |
| *neuroligin 1 (nlg1)* | vdrc 42616 | postsynaptic cell surface protein, cell adhesion, excitatory synapses | | BG380, Dicer2 | no difference from UAS-control (luciferase) RNAi |
| *defective proboscis extension response 4 (dpr4)* | vdrc 28519 | cell surface receptor for DIPs, adhesion protein, synapse organization | | BG380, Dicer2 | no difference from UAS-control (luciferase) RNAi |
| | vdrc 102905 | | | BG380, Dicer2 | no difference from UAS-control (luciferase) RNAi |
| | vdrc 28518 | | | BG380, Dicer2 | no difference from UAS-control (luciferase) RNAi |
| *fasciculin II (fasII)* | RRID:BDSC_34084 | cell adhesion molecule, axonal pathfinding/guidance | | BG380, Dicer2 | no difference from UAS-control (luciferase) RNAi |
| | vdrc 8393 | | | BG380, Dicer2 | no difference from UAS-control (luciferase) RNAi |
| | vdrc 36350 | | | BG380, Dicer2 | no difference from UAS-control (luciferase) RNAi |
| *comatose/NSF1 (comt)* | RRID:BDSC_27890 | N-ethylmaleimide-sensitive factor 1, role disassembly of SNARE complexes | | D42 | no difference from UAS-control (luciferase) RNAi |
| *semaphorin 2b (sema2b)* | RRID:BDSC_28932 | secreted semaphorin, involed in embryonic axonal guidance | | BG380, Dicer2 | no difference from UAS-control (luciferase) RNAi |
| *plexin B (plexB)* | RRID:BDSC_57813 | Acts as a receptor for Sema-2a and seems to suppress axon branching | | BG380, Dicer2 | no difference from UAS-control (luciferase) RNAi |
| *neuroglian (Nrg)* | RRID:BDSC_37496 | L1-type cell adhesion molecule | | BG380, Dicer2 | no difference from UAS-control (luciferase) RNAi |
| *wishful thinking (wit)* | RRID:BDSC_25949 | BMP type II receptor with known functions synaptic development and maintenance at the Drosophila NMJ | | BG380, Dicer2 | no effect on *puc*-lacZ induction in uninjured animals. *puc*-lacZ induction 24 post peripheral nerve crush is similar to wild type animals. |

| | | | | | |
|---|---|---|---|---|---|
| | vdrc 103808 | | | BG380, Dicer2 | " |
| *thickveins (tkv)* | RRID:BDSC_41904 | BMP type I receptor with known functions synaptic development and maintenance at the Drosophila NMJ | | BG380, Dicer2 | no effect on *puc*-lacZ induction in uninjured animals. *puc*-lacZ induction 24 post peripheral nerve crush is similar to wild type animals. |
